# Antibiotics at Clinical Concentrations Show Limited Effectivity Against Acute and Chronic Intracellular *S. aureus* Infections in Osteocytes

**DOI:** 10.1101/2023.12.17.572089

**Authors:** Anja R. Zelmer, Dongqing Yang, Nicholas J. Gunn, L. Bogdan Solomon, Renjy Nelson, Stephen P. Kidd, Katharina Richter, Gerald J. Atkins

## Abstract

**Objectives:** Case numbers of osteomyelitis are rising and chronic infections remain difficult to cure. While it is known that the major pathogen *Staphylococcus aureus* can persist intracellularly in osteocytes, the effectivity of antibiotics against this condition remains largely unknown. We sought to determine if current clinically utilised antibiotics were capable of clearing an intracellular osteocyte *S. aureus* infection.

**Methods:** Rifampicin, vancomycin, levofloxacin, ofloxacin, amoxicillin, oxacillin, doxycycline, linezolid, gentamicin and tigecycline were assessed for their MIC and minimum bactericidal concentrations (MBC) against 11 *S. aureus* clinical isolates and the reference strain ATCC 25923, at pH 5.0 and 7.2 to mimic lysosomal and cytoplasmic environments, respectively. Those antibiotics whose bone achievable concentration was commonly above their respective MICs for the strains tested were further assayed in a human osteocyte infection model under either acute or chronic conditions. Osteocyte-like cells were treated at 1, 4 and 10x the MIC for 1 and 7 days following infection (acute model), or after 14 days of infection (chronic model). The intracellular effectivity of each antibiotic was measured in terms of colony forming unit (CFU) reduction, small colony variant (SCV) formation and bacterial mRNA expression change.

**Results:** Only rifampicin, levofloxacin and linezolid reduced intracellular CFU numbers significantly in the acute model. The effect was larger after 7 days compared to 1 day of treatment. However, no treatment reduced the quantity of bacterial mRNA, nor prevented non-culturable bacteria from returning to a culturable state.

**Discussion:** These findings indicate that *S. aureus* adapts phenotypically during intracellular infection of osteocytes, adopting a reversible quiescent state which is protected against antibiotics, even at 10x their MIC. Thus, new therapeutic approaches are necessary to cure *S. aureus* intracellular infections in osteocytes.

## Introduction

Osteomyelitis is challenging to treat, and recurrent and chronic infections associated with treatment failure occur in up to 46% of cases [1, 2]. The most prevalent pathogen, *Staphylococcus aureus*, also has the highest treatment failure rate [2–4], and potentially contributing to this, *S. aureus* can persist intracellularly in phagosomes or the cytoplasm [5, 6] in immune cells and bone cells, including osteocytes [7].

The intracellular effectivity of antibiotics is difficult to predict and is often reduced due to host factors including intracellular metabolism, drug efflux, the low pH of certain intracellular compartments, as well as potential off-target effects on the pathogen clearance mechanisms of autophagy/xenophagy [8–11]. Bacterial factors also play a role, for example chronic osteomyelitis is associated with increased small colony variant (SCV) formation [4, 12, 13], resulting in higher antibiotic tolerance and immune protection [14, 15]. Furthermore, persister cells, a subpopulation of dormant bacteria with reduced metabolism, also confer antibiotic insensitivity [16]. Thus, to treat chronic osteomyelitis effectively, both intracellular location and phenotypic adaptation are potential factors to consider.

Osteocytes form a syncytial network throughout the bone and are the most abundant and long-lived resident population [17], theoretically providing an ideal niche for long-term infections, as has been reported with phenotypic adaption into SCV [12, 18, 19].

Here, we investigated the effectivity of clinically used antibiotics at concentrations predicted to be achievable in the bone in a human osteocyte-like intracellular infection model against *S. aureus*. Acute and chronic infections, short and long-term antibiotic treatments with antibiotics of different classes and mechanisms of action, with differing levels of evidence for effectivity [20], were investigated.

## Methods

### Osteocyte-like Culture

SaOS-2 cells (American Type Culture Collection (ATCC), Manassas, USA) were passaged and differentiated to an osteocyte-like stage, as previously described [21]. The osteocyte-like phenotype of the resulting cultures was confirmed by increased expression of phenotypic markers *COL1A1*, *RUNX2*, *MEPE*, *BGLAP, DMP1* and *SOST* and *in vitro* mineralisation (**Supp. Fig. 1**) [21].

### Bacterial Isolates and Growth Conditions

Institutional human research ethics clearance for this study was obtained (CALHN Approval 16633). Eleven *S. aureus* clinical isolates (CI) from the soft tissue wound or bone from patients undergoing treatment for diabetic foot infection and the strain ATCC25923 were used. Bacteria were grown in terrific broth (TB; Invitrogen, Waltham, USA), as previously described [21]. Viable bacterial concentrations were estimated from a CFU/ml *vs* OD_630 nm_ standard curve, then validated by plating dilutions on nutrient broth (NB)-agar (Chem-supply, Gillman, Australia).

### MIC and Minimal Bactericidal Concentration (MBC)

MICs for tigecycline, linezolid, oxacillin-sodium-salt-monohydrate, levofloxacin, ofloxacin (Glentham Life Science, Corsham, UK), doxycycline-hydrate, vancomycin, amoxicillin and rifampicin (Sigma-Aldrich, St Louis, USA) were determined with the micro-dilution method in TB at pH 5.0 and pH 7.2 [22]. For MBC evaluation all clear wells and the first well showing turbidity were spot-plated and colony growth was observed after 24 and 48 h. European Committee on Antimicrobial Susceptibility Testing (EUCAST)-tables [23] were used to define resistance.

### Intracellular Antibacterial Activity

CI 8 was grown as described above and diluted to a multiplicity of infection (MOI) of 50 in phosphate buffered saline (PBS). Intracellular infection was performed, as previously described [19, 24]. Briefly, differentiated SaOS2 cells were exposed to the bacterial suspension or sterile PBS for 2h. Following washing with PBS, all cells were re-incubated with 10 mg/l lysostaphin for 2 h in antibiotic-free media and then treated with antibiotics at 1x, 4x, 10x their respective MIC (i.e., 0.25, 1, 2.5 mg/l doxycycline-hydrate, 0.016, 0.064, 0.16 mg/l rifampicin, 0.0625, 0.25, 0.625 mg/l oxacillin, 1, 4, 10 mg/l levofloxacin and 2, 8, 20 mg/l linezolid) either immediately or after 14 days. Cells were harvested for analysis after 1, 7, 15 and 21 days. To test for outgrowth after antibiotic treatment in the chronic model, the 10x MIC treatments were removed and the cells washed with PBS, before changing to an antibiotic- and lysostaphin-free media. Media were spot-plated regularly and observed for changes in colour and cloudiness daily. After 11 days, cells were washed and lysed to identify any intracellular CFU that did not grow into the media. For CFU recovery, culture supernatants were tested for viable bacteria by spot-plating. The host-cells were lysed in sterile water for 20 min, 1:10 serially diluted and plated on NB-agar.

### Measurement of Gene Expression

Cell supernatant was removed, and RNA isolated using TRI Reagent (Merck Life Science, VIC, Australia). After DNAse digestion, complementary DNA was synthesised with the iScript gDNA clear cDNA Synthesis kit (BioRad, Hercules, USA), all as per manufacturer’s instructions. DNA was extracted with the Viagen DirectPCR® DNA Extraction System^TM^ and 1% proteinase K (Viagen Biotech Inc, CA, USA) for 10 min at room temperature, 30 min at 55°C and 15 min at 85°C. DNA was digested with Ncol-HF (New England BioLabs, Notting Hill, Australia). Real-time PCR reactions were performed using RT2 Forget-Me-Not^TM^ EvaGreen qPCR Mastermix (Biotium, CA, USA) on a CFX Connect Real-Time-PCR-System (BioRad). Measurement of effects on bacterial housekeeping gene levels in response to a 10x MIC dose of each antibiotic was made against infected but untreated control or the corresponding 1x MIC treatment. Oligonucleotide primer sequences are shown in **Supplementary Table 1**.

### Statistics

Differences in CFU, gene expression and proportion of SCV (%SCV) were analysed with either a 1- or 2-way ANOVA with a Tuckey’s multiple comparison or an unpaired t -Test (α = 0.05) using GraphPad Prism version 10.2.3 (GraphPad Software, La Jolla, USA). Graphs herein show mean values ± standard deviation (SD).

## Results

### MIC and MBC determination

To determine which antibiotics could be effective at concentrations achievable in bone, we determined their respective MIC and MBC against 11 CIs of *S. aureus* and ATCC 25923 at both pH 5.0 and pH 7.2 (**Table 1**). All CIs were susceptible to vancomycin, linezolid, gentamicin and tigecycline, while five strains were resistant to levofloxacin, ofloxacin and oxacillin. The MBCs for some strains were at least 4-fold higher than the MICs for rifampicin, oxacillin, doxycycline, linezolid and tigecycline, indicating a strong bacteriostatic effect. Some antibiotics showed pH dependency, with levofloxacin, ofloxacin, doxycycline and tigecycline being at least 4-times more effective at pH 7.2 than at pH 5.0 and rifampicin, oxacillin and gentamicin being more effective at pH 5.0 than pH 7.2, consistent with previous studies [20].

**Table 1:**
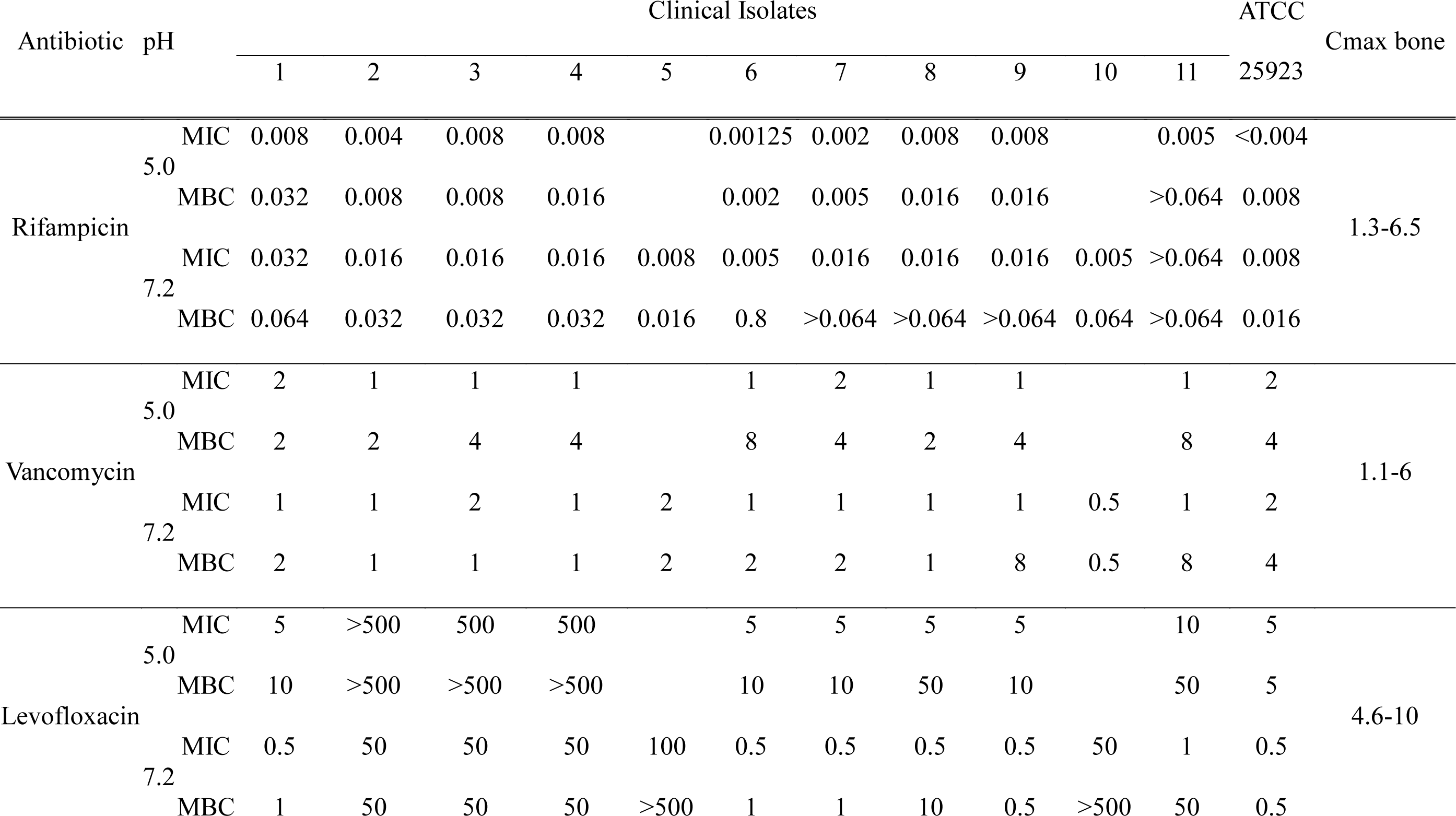
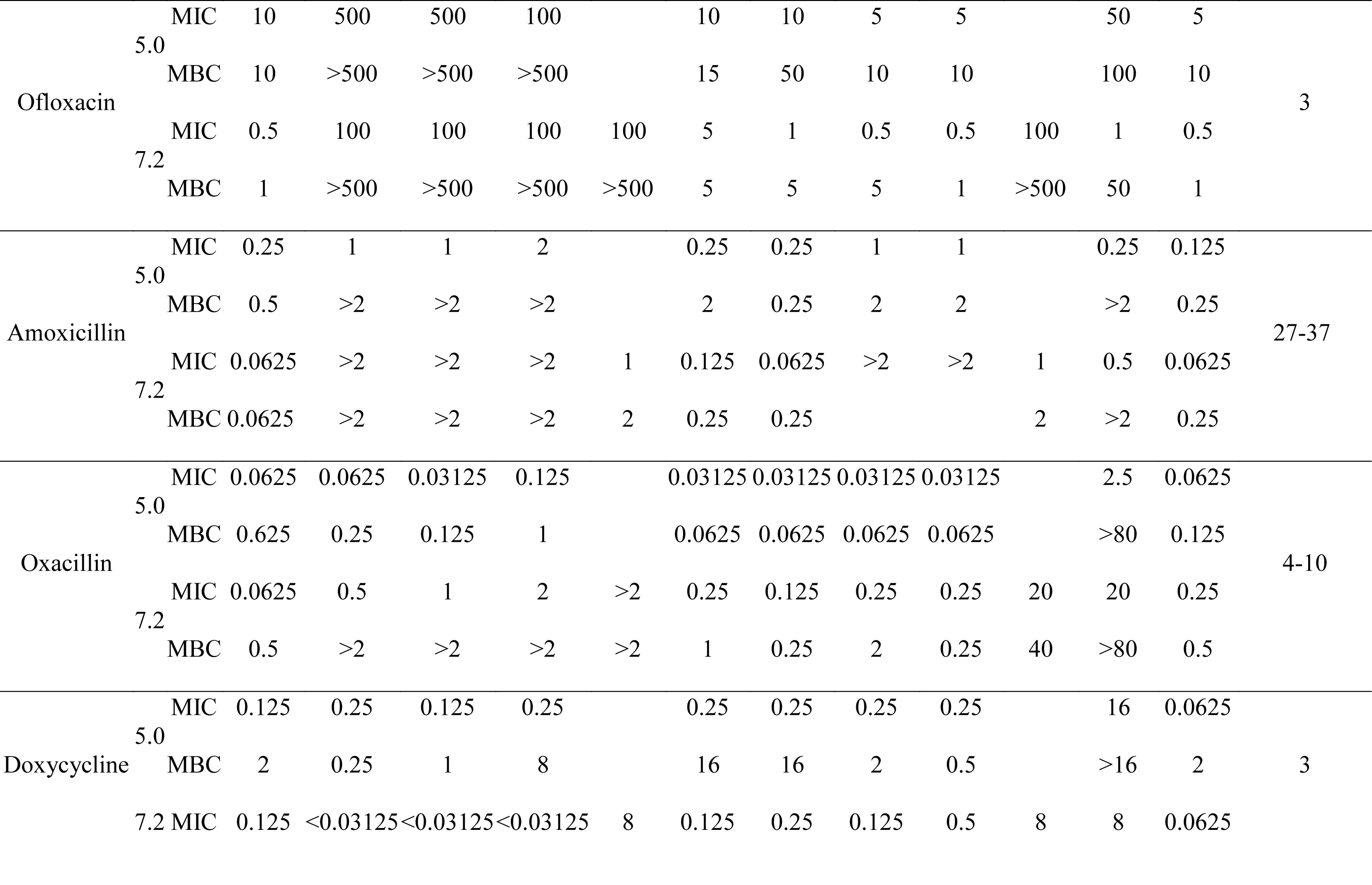
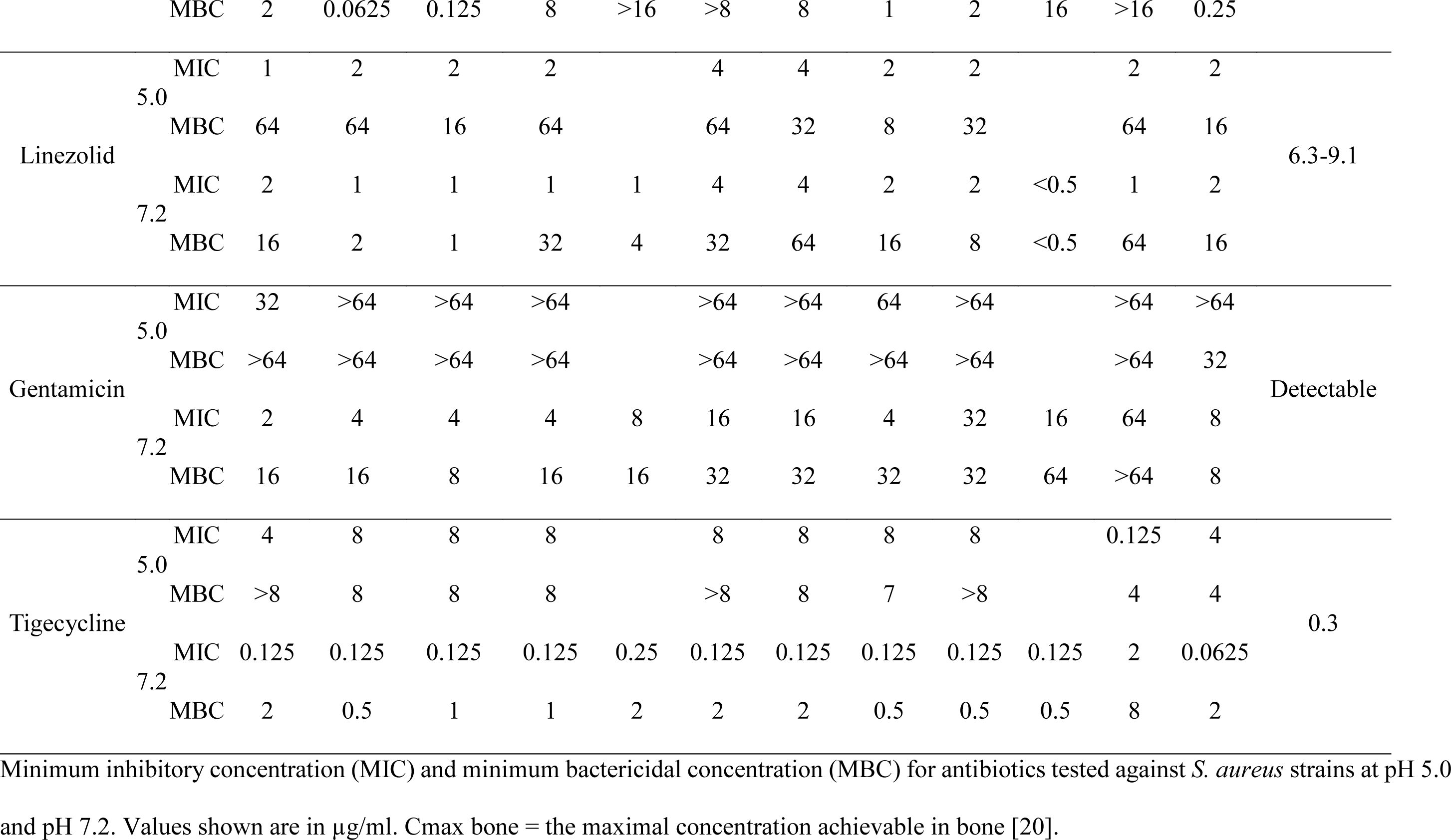
Antimicrobial susceptibility of *S. aureus* and antimicrobial concentrations.

The MIC and MBC values for each strain were compared to respective bone-achievable concentrations [20]. Antibiotics that could reach a potentially therapeutic concentration in at least 6 strains (i.e. >4x MIC [20]) were further tested in an intracellular infection model: doxycycline, oxacillin, levofloxacin, rifampicin and linezolid.

### Infected Host-cell Survival

To ensure that neither the bacterial infection itself, nor the antibiotic treatments affected host-cell viability, we conducted live/dead staining at each time point and measured the *18S* rRNA expression. There was no significant change due to the infection or any treatment at any time point for either measure, as shown for levofloxacin (**Supp. Fig. 2**).

### Intracellular Antibacterial Activity

We tested doxycycline, oxacillin, levofloxacin, rifampicin and linezolid for intracellular antibacterial activity at 1, 4 and 10x their respective MIC for either 1 or 7 days and started either immediately (acute model) or 15 days (chronic model) post-infection (**Fig. 1A**).

**Figure 1:**
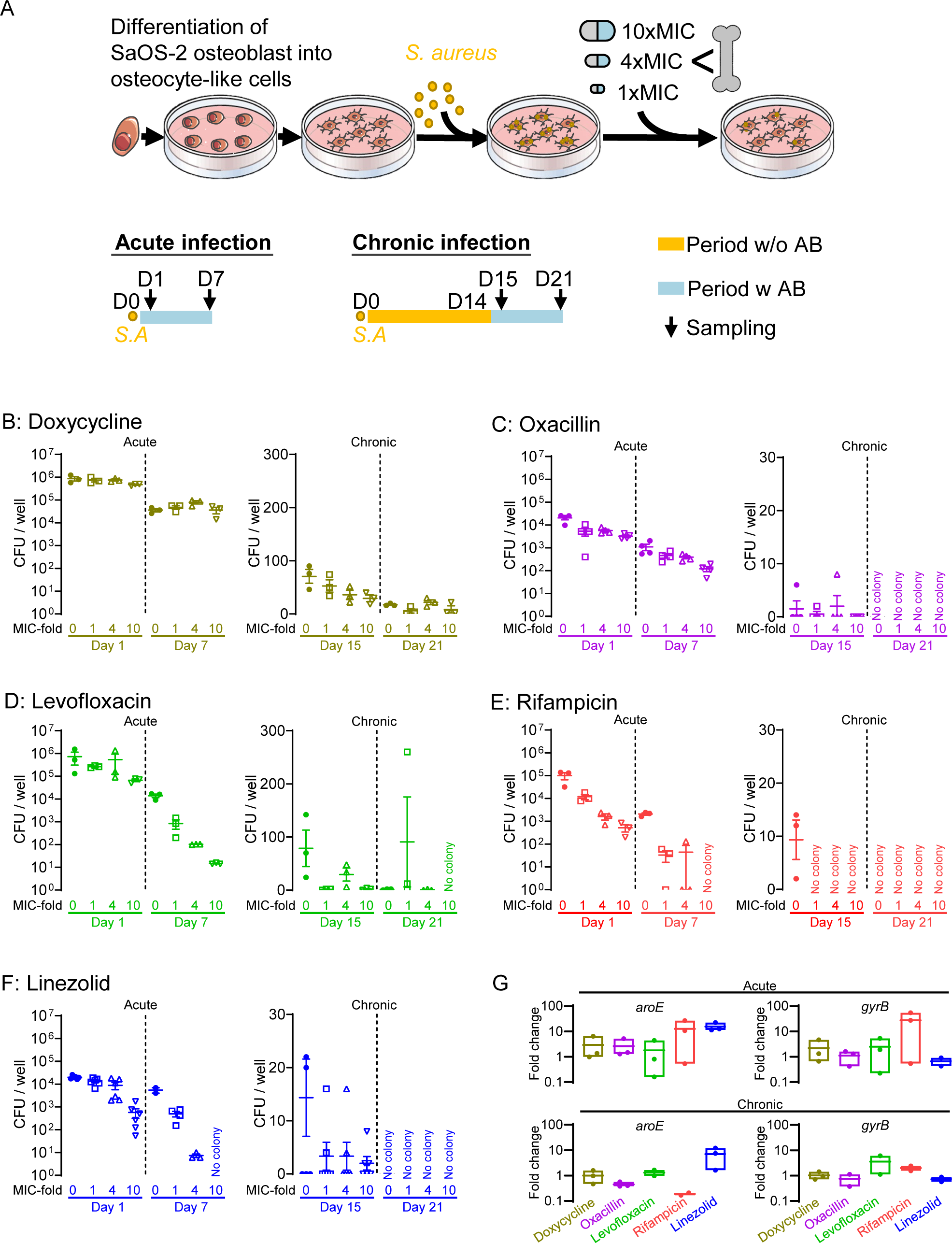
Intracellular bacterial culturability in acute and chronic *S. aureus* infection models. A) Schematic of the experimental design of the osteocyte infection and antibiotic treatment model. Upon the introduction of *S. aureus* infection to SaOS2-OY cells, antibiotic (AB) treatments were provided to cultures either immediately after the infection on day 0 for the acute model or 14 days post-infection for the chronic model. Recovered CFU from acute (sampled at days 1 and 7) and chronic (sampled at days 15 and 21) models under antibiotic treatments of B) doxycycline, C) oxacillin, D) levofloxacin, E) rifampicin and F) linezolid, at the doses of 0, 1, 4 and 10x the respective MIC; G) Fold-change in mRNA levels of two bacterial genes, *aroE* and *gyrB*, among individual antibiotic treatment groups compared to expression in the untreated control, on days 7 and 21 for acute and chronic models, respectively.

For the acute model, CFU recovery declined between 1 and 7 days post-infection for all conditions (**Fig. 1B-F**). Beyond this, doxycycline had no effect on the number of intracellular CFUs (**Fig. 1B**, **Table 2**). Oxacillin caused a significant, but clinically irrelevant CFU reduction after 1 day of treatment at all concentrations, with no change after 7 days (**Fig. 1C**, **Table 2**). Levofloxacin was ineffective after 1 day, however, 7 days of treatment resulted in a significant and clinically relevant intracellular CFU-reduction of up to 2.98-log (**Fig. 1D, Table 2**). Rifampicin caused the highest significant dose-dependent CFU reduction after 1 day of up to 2.28-log, which increased up to 3.32-log after 7 days of treatment (**Fig. 1E, Table 2**). The highest significant CFU reduction of 3.74-log was achieved after 7 days of treatment with linezolid, however, linezolid had a significant but smaller impact after 1 day of treatment of 1.54-log (**Fig. 1F, Table 2**).

**Table 2:**
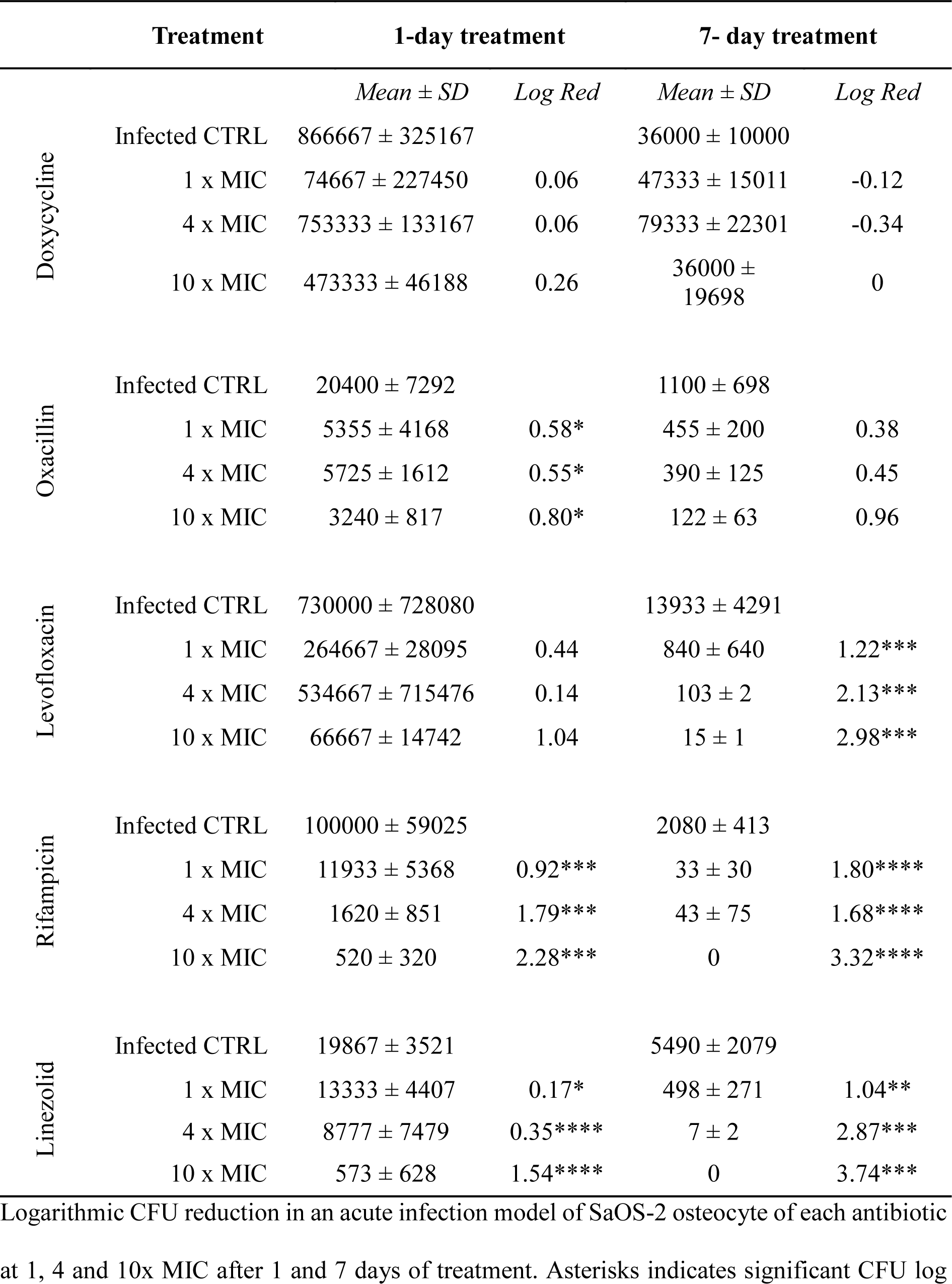
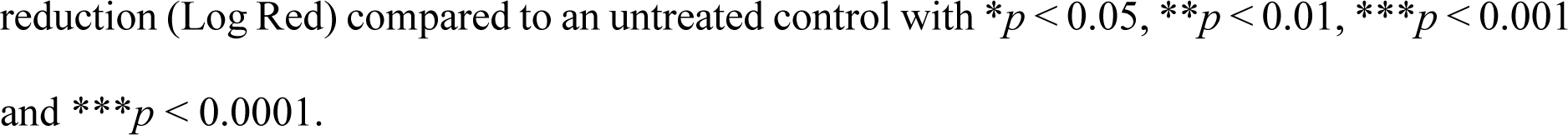
Antimicrobial effect on colony forming units (CFU) in an acute intra-osteocytic *S. aureus* infection.

| Treatment |  | 1-day treatment |  | 7- day treatment |  |
| --- | --- | --- | --- | --- | --- |
|  |  | *Mean ± SD* | *Log Red* | *Mean ± SD* | *Log Red* |
| Doxycycline | Infected CTRL | 866667 ± 325167 |  | 36000 ± 10000 |  |
|  | 1 x MIC | 74667 ± 227450 | 0.06 | 47333 ± 15011 | -0.12 |
|  | 4 x MIC | 753333 ± 133167 | 0.06 | 79333 ± 22301 | -0.34 |
|  | 10 x MIC | 473333 ± 46188 | 0.26 | 36000 ± 19698 | 0 |
| Oxacillin | Infected CTRL | 20400 ± 7292 |  | 1100 ± 698 |  |
|  | 1 x MIC | 5355 ± 4168 | 0.58* | 455 ± 200 | 0.38 |
|  | 4 x MIC | 5725 ± 1612 | 0.55* | 390 ± 125 | 0.45 |
|  | 10 x MIC | 3240 ± 817 | 0.80* | 122 ± 63 | 0.96 |
| Levofloxacin | Infected CTRL | 730000 ± 728080 |  | 13933 ± 4291 |  |
|  | 1 x MIC | 264667 ± 28095 | 0.44 | 840 ± 640 | 1.22*** |
|  | 4 x MIC | 534667 ± 715476 | 0.14 | 103 ± 2 | 2.13*** |
|  | 10 x MIC | 66667 ± 14742 | 1.04 | 15 ± 1 | 2.98*** |
| Rifampicin | Infected CTRL | 100000 ± 59025 |  | 2080 ± 413 |  |
|  | 1 x MIC | 11933 ± 5368 | 0.92*** | 33 ± 30 | 1.80**** |
|  | 4 x MIC | 1620 ± 851 | 1.79*** | 43 ± 75 | 1.68**** |
|  | 10 x MIC | 520 ± 320 | 2.28*** | 0 | 3.32**** |
| Linezolid | Infected CTRL | 19867 ± 3521 |  | 5490 ± 2079 |  |
|  | 1 x MIC | 13333 ± 4407 | 0.17* | 498 ± 271 | 1.04** |
|  | 4 x MIC | 8777 ± 7479 | 0.35**** | 7 ± 2 | 2.87*** |
|  | 10 x MIC | 573 ± 628 | 1.54**** | 0 | 3.74*** |

Logarithmic CFU reduction in an acute infection model of SaOS-2 osteocyte of each antibiotic at 1, 4 and 10x MIC after 1 and 7 days of treatment. Asterisks indicates significant CFU log
reduction (Log Red) compared to an untreated control with $*p < 0.05$ , $**p < 0.01$ , $***p < 0.001$ and $***p < 0.0001$ .

In the chronic infection model, few if any CFU were recovered, such that the change in CFU number could not be statistically analysed (**Fig. 1B-F**).

To measure the effects of treatments on bacterial metabolism independent of effects on culturability, we also examined bacterial mRNA expression, using *aroE* and *gyrB* as representative genes. For the acute model, the impact of antibiotic treatments on bacterial mRNA levels were not concordant with CFU recovery. Even at 10x MIC for 7 days no antibiotic treatment reduced intracellular mRNA levels compared to the untreated control (**Fig. 1G**). Bacterial mRNA expression in the chronic model was similar to that of the acute model and did not significantly change with any treatment throughout the time course (**Fig. 1G**).

To further examine potential effects of antibiotics on growth phenotype, the percentage of SCV was recorded. Doxycycline, oxacillin, rifampicin and linezolid did not significantly impact the %SCV (**Fig. 2B, C, E, F**). Levofloxacin caused a dose-dependent increase in the %SCV up to 28.21% (**Fig. 2D**).

**Figure 2:**
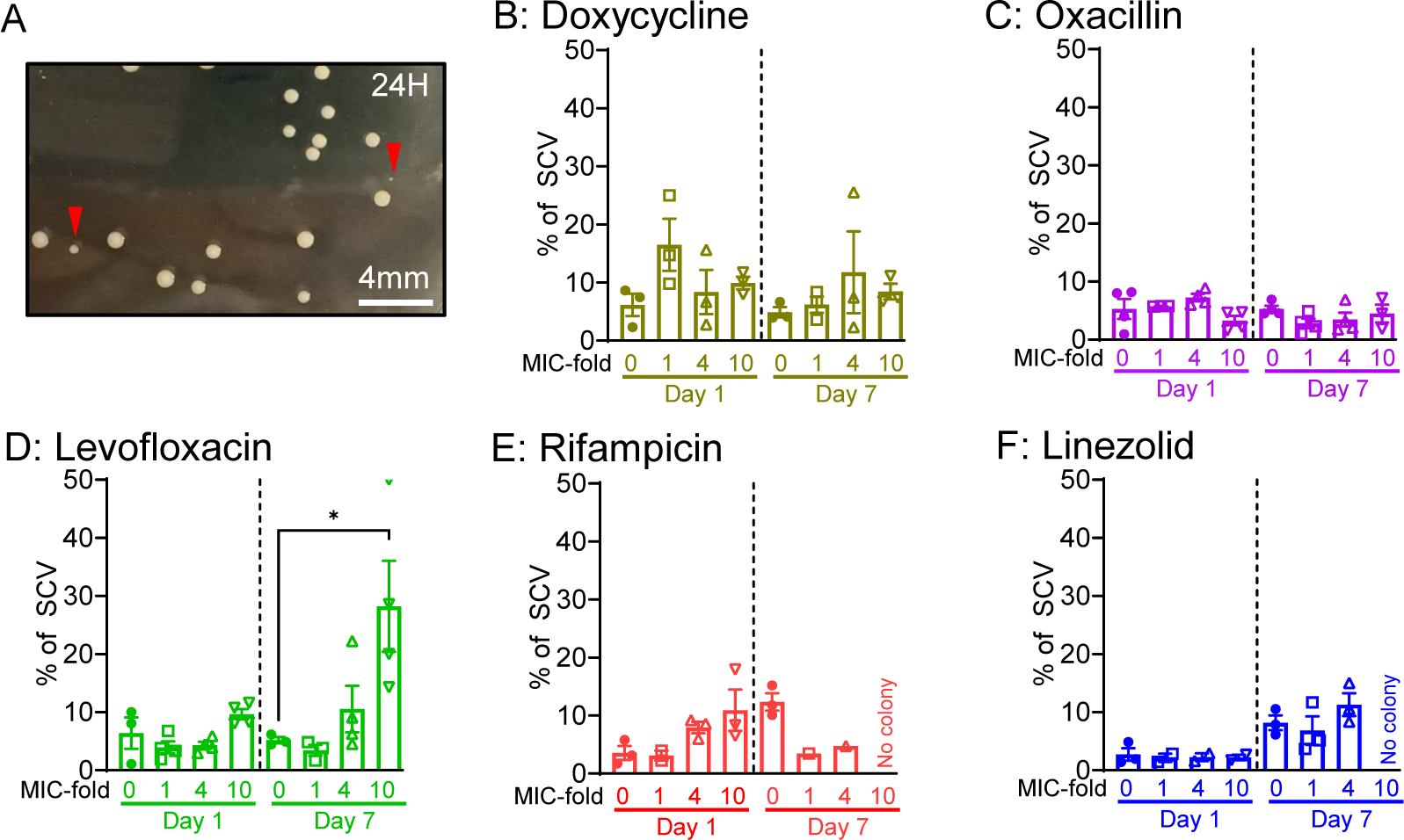
Effects of antibiotics on colony phenotype of *S. aureus* recovered from an intracellular infection in osteocytes. A) Example image showing small colony variants (SCV; *red arrows*), defined here as a colony after 24 hours of culture on nutrient agar measuring <0.5 mm; B-F) The %SCV from host-cell lysates was determined after 1 and 7 days of treatment with B) doxycycline, C) oxacillin, D) levofloxacin, E) rifampicin and F) linezolid, at 1x, 4x and 10x their respective MIC (*n* ≥ 3). Significant differences are indicated by * *p* < 0.05.

### Bacterial Resuscitation

To test the ability of antibiotic-treated cultures displaying non-culturable infections to give rise to a culturable infection, a bacterial outgrowth assay was performed. Following robust treatment of intracellular bacteria with a 3 day 10x MIC antibiotic regimen, antibiotic pressure was removed. As shown in **Figure 3**, bacterial resuscitation occurred in untreated control cultures after 7 days, with approximately 60% of wells showing outgrowth. Outgrowth occurrence was similar for doxycycline, levofloxacin and linezolid, while outgrowth was evident earlier, at 4 days, for oxacillin-treated cultures. Most wells containing culturable extracellular bacteria also had culturable bacteria in the host cell lysate at the experimental end. However, rifampicin treatment only yielded culturable bacteria from the host cell lysate (**Fig. 3**).

**Figure 3:**
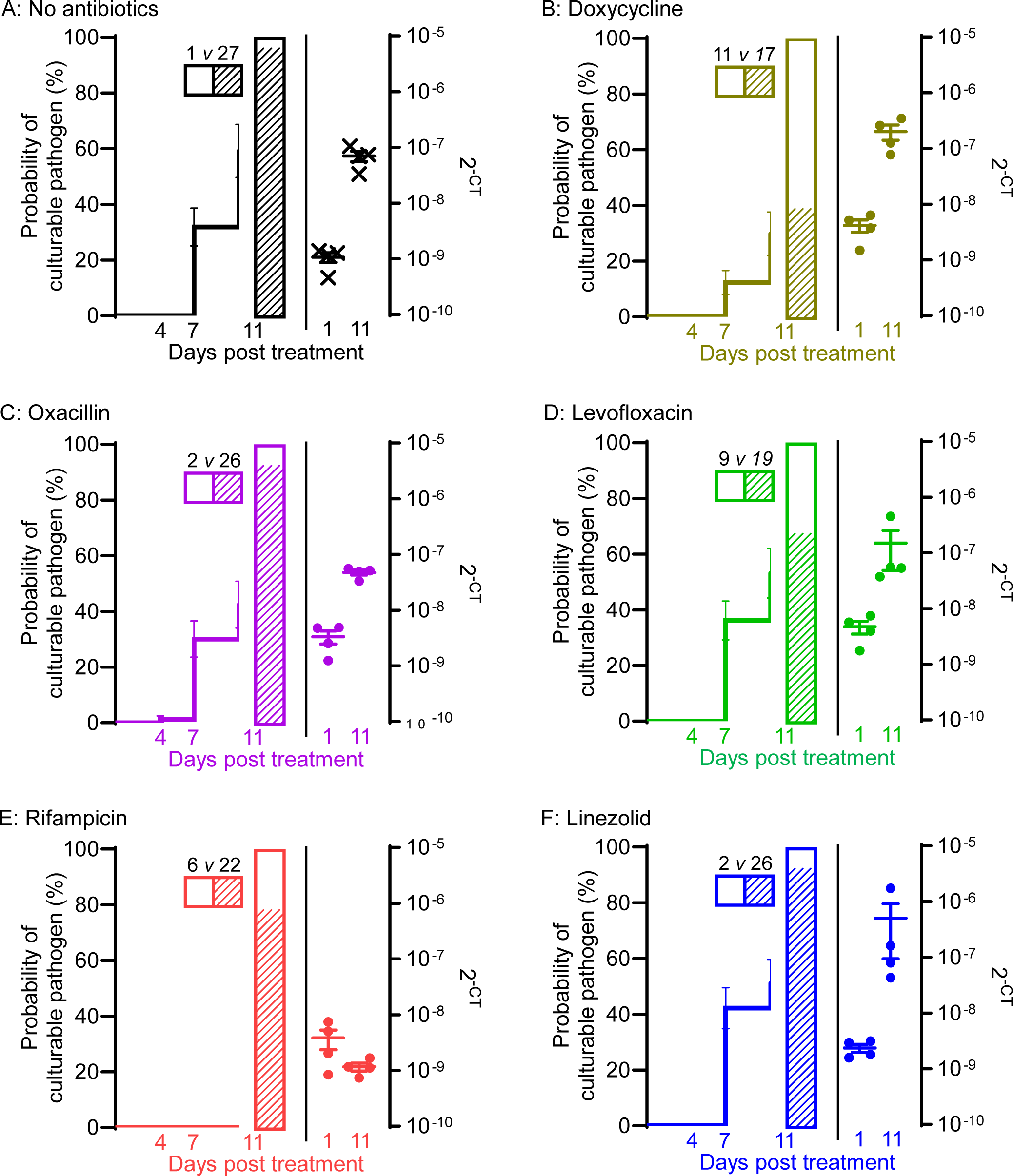
Outgrowth of intracellular *S. aureus* from a non-culturable state. SaOS2-OY cells were infected with *S. aureus* and intracellular bacteria *were* confirmed non-culturable by 14 days post-infection. All cells were then treated for 3 days with either antibiotic-free media or with antibiotics at their respective 10x MIC. For all treatment groups (A: no antibiotic; B: doxycycline; C: oxacillin; D: levofloxacin; E: Rifampicin; F: Linezolid), cell culture media were tested for extracellular bacteria culturability from potential outgrowth on days 4, 7 and 11 (n = 32 wells per condition, line graphs on left Y-axis). Cell lysates from the experiment endpoint (day 11) were tested for intracellular bacterial culturability (n = 28 wells per condition, bar graphs on left Y-axis, hatched pattern indicates growth). The number of wells with or without outgrowth are shown adjacent. Total DNA isolated from cell lysates on days 1 and 11 was measured for intracellular bacterial genome levels by qPCR (n = 4 wells per condition; dot graphs on right Y-axis).

Intracellular bacterial DNA levels tended to mirror the change in bacterial outgrowth, clearly increasing between days 1-11 post-treatment, consistent with increased bacterial replication, with the exception of rifampicin-treated cultures which displayed stable bacterial levels (**Fig. 3**).

## Discussion

This study highlights an important limitation of currently available antibiotics, in that despite bone penetrance, while culturability may be impaired, intracellular bacterial viability and activity (DNA and mRNA levels) persist, indicating a switch to a non-culturable but nevertheless potentially virulent phenotype. This serves as further evidence that phenotype switching to low- or non-culturable phenotypes is an important and emerging aspect of intracellular infections in osteomyelitis and is likely to contribute to antimicrobial treatment failure and persistent infection [11, 12, 25].

Rifampicin is among the most demonstrably effective antibiotics against intracellular *S. aureus* [20]. This was confirmed here as it demonstrated a 2-8-fold higher sensitivity at pH 5.0 compared to pH 7.2, indicating high lysosomal activity, and up to a 3.7-log CFU reduction in the osteocyte model. Furthermore, rifampicin was the only treatment that did not yield culturable bacteria in the media after antibiotic removal and the only antibiotic that inhibited bacterial population growth following the cessation of treatment, as evidenced by the lack of increase in bacterial DNA levels, at least within the time frame of the analysis.

Linezolid was the only protein-biosynthesis-inhibitor tested that effectively reduced intracellular CFU numbers up to a 3.74-log, in particular after a 7 day treatment. However, it appeared to impact the growth phenotype, as apparent persister cells readily reverted and escaped after antibiotic removal.

The fluoroquinolones ofloxacin and levofloxacin showed an approximately 10-fold reduced susceptibility at pH 5.0 and six CIs were resistant [23]. This resulted in a higher MIC than the 3 mg/l achievable in bone for ofloxacin, however, for levofloxacin, when susceptible, the MIC was between 0.5-5 mg/l at both pH levels tested, a range that easily falls below the maximum concentration of 10 mg/l achievable in bone [20]. Levofloxacin reduced CFU numbers up to 2.98-log after 7 days of treatment. Still, levofloxacin failed to reduce the bacterial mRNA levels at any stage and showed a significant dose-dependent increase in %SCV after 7 days of treatment. Therefore, fluoroquinolones might be effective in reducing intracellular CFU numbers if the bacterial strain is sensitive, however, they also might induce phenotypic switches to less antibiotic and immune-susceptible phenotypes.

Doxycycline showed a strongly strain-dependent MIC, lower than the reported maximum bone concentration up to 3 mg/l. Nevertheless, it proved to be ineffective in the osteocyte model.

The three cell wall inhibitors (oxacillin, amoxicillin and vancomycin) were found to be ineffective in reducing intracellular bacterial numbers. Oxacillin was much more effective at pH 5.0 than at pH 7.2 and at a much lower concentration than the maximum bone level of 4- 10 mg/l [20]; however, this was ineffective in the osteocyte model. Gentamicin is often described as only being effective against extracellular pathogens [26]. Nonetheless, the cell:plasma ratio measured in macrophages between 4.4 and 6.8 is comparable to other antibiotics with intracellular activity, such as linezolid and oxacillin [8, 9]. We observed a 2- to 8-fold reduced effectivity at pH 5.0 compared to pH 7.2, which might contribute to the low intracellular activity of 1-3-log reported in bone cells [13, 20], when given at concentrations that exceed those that can be expected in the bone due to poor bone penetration [20]. Therefore, gentamicin does not seem to be a suitable candidate to treat chronic osteomyelitis. Tigecycline and vancomycin showed higher MICs than can be achieved in the bone and were therefore not tested in the osteocyte model.

Limitations of this study include the number of antibiotics tested and the mono-therapeutic approach. Clinically utilised antibiotics were selected to represent various mechanisms of action and included those that were predicted to be most effective [20]. Further, even though the MIC/MBC were tested in multiple CIs, for feasibility purposes, the intracellular infection assay was only conducted with one strain. However, this isolate, an actual osteomyelitis strain, brings further clinical relevance to the findings. The purpose of the model utilised was to facilitate high-throughput testing of antimicrobial effectivity against intracellular *S. aureus* in a human osteocyte-like context. To achieve reproducibility, we used an osteocyte-like cell line model rather than primary cells [12], which might behave differently. However, osteogenically differentiated SaOS-2 is a validated human osteocyte-like model for infection experiments, shown to possess key properties of primary cells [19, 21, 27], as also verified here.

In conclusion, the most effective antibiotics at clinically-relevant concentrations for reducing intracellular CFUs in a human osteocyte-like cell model were rifampicin, levofloxacin and linezolid. Treatment duration has a significant impact on the effectivity of these antibiotics. However, treatments that were effective in reducing bacterial CFU numbers, did not eradicate the bacteria, as intracellular bacterial DNA and mRNA remained detectable and bacteria may switch to a non-culturable state and regrow once antibiotic pressure is removed. In this respect, antibiotics treatments serve as agents for selecting persisting bacterial populations, rather than curing the intracellular infection. The impact of unsuccessful antimicrobial treatment of intracellular infection on the induction of antibiotic resistance is yet to be investigated. The model described herein could be used to examine strain-specific antimicrobial effectivity for the purposes of optimising treatment of osteomyelitis, and to test new or emerging drugs in this context.

## Acknowledgments

This work was supported by funding from the National Health and Medical Research Council of Australia (NHMRC; Grant No. 2011042) awarded to G.J.A. and D.Y.. A.R.Z. and N.J.G. were supported by University of Adelaide Faculty of Health and Medical Sciences Postgraduate Research Scholarships. The authors wish to thank Mr. James Lee and Professor Robert Fitridge (The University of Adelaide) for providing the clinical *S. aureus* isolates for this study.

## Author Contributions

ARZ project conception, performed experiments, manuscript draft

DY, NJG, KR, SPK experimental support, manuscript development

LBG, RN manuscript development

GJA, project conception, manuscript development, communicating author.

---

**Supplementary Figure 1:** Osteocytic differentiation of SaOS-2 cells. A-F) Osteogenic gene expression of osteocyte markers during differentiation relative to *18S* rRNA levels (*n* = 4). \**p* < 0.05, \*\*\**p* < 0.001 and \*\*\**p* < 0.0001. Gene expression was normalised to that of the housekeeping gene *18S* and calculated with the 2^−ΔCt^ method. G) Von Kossa stain for phosphate and Alizarin Red stain for calcium to observe mineralisation of osteocytes after 7 and 14 days of differentiation (*n* = 4).

**Supplementary Figure 2:** Survival of host-cells during intracellular infection. A) Levels of *18S* rRNA over time in infected control, uninfected control or treatment with levofloxacin (as a representative example for all treatments; *n* = 4). B) Live/dead staining of infected and uninfected controls. Cells were cultured as described above, on Cell Imaging Plates (Eppendorf, Germany). Immediately before the infection or 1, 7, 15 and 21 days after the infection the media of infected and uninfected wells were incubated in the dark at 37°C/ 5% CO_2_/ 1% O_2_ with eBioscience TM Calcein Violet 450 AM Viability Dye (live, Invitrogen) and Ethidium Homodimer III (dead, Biotium) for 30 min. Confocal images were then taken with an Olympus FV3000 confocal microscope (Olympus, Tokyo, Japan) and processed with ImageJ software (Bethesda, USA) to obtain relative intensities of live/dead stained cells. Data shown as means ± standard error of the mean for 4 biological replicates, in at least 3 regions of interest per well.

**Supplementary Table 1:**
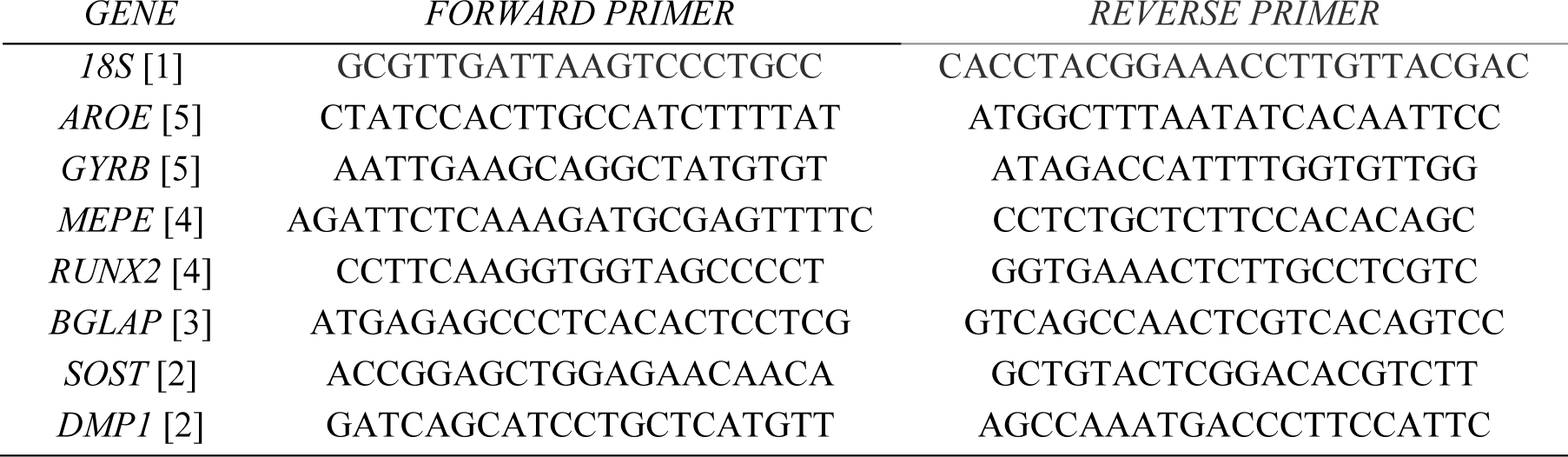
Oligonucleotide primer sequences for real time PCR.

