## Supplementary material for "Antibiotics at Clinical Concentrations Show Limited Effectivity Against Acute and Chronic Intracellular *S. aureus* Infections in Osteocytes": Suppl Fig 1

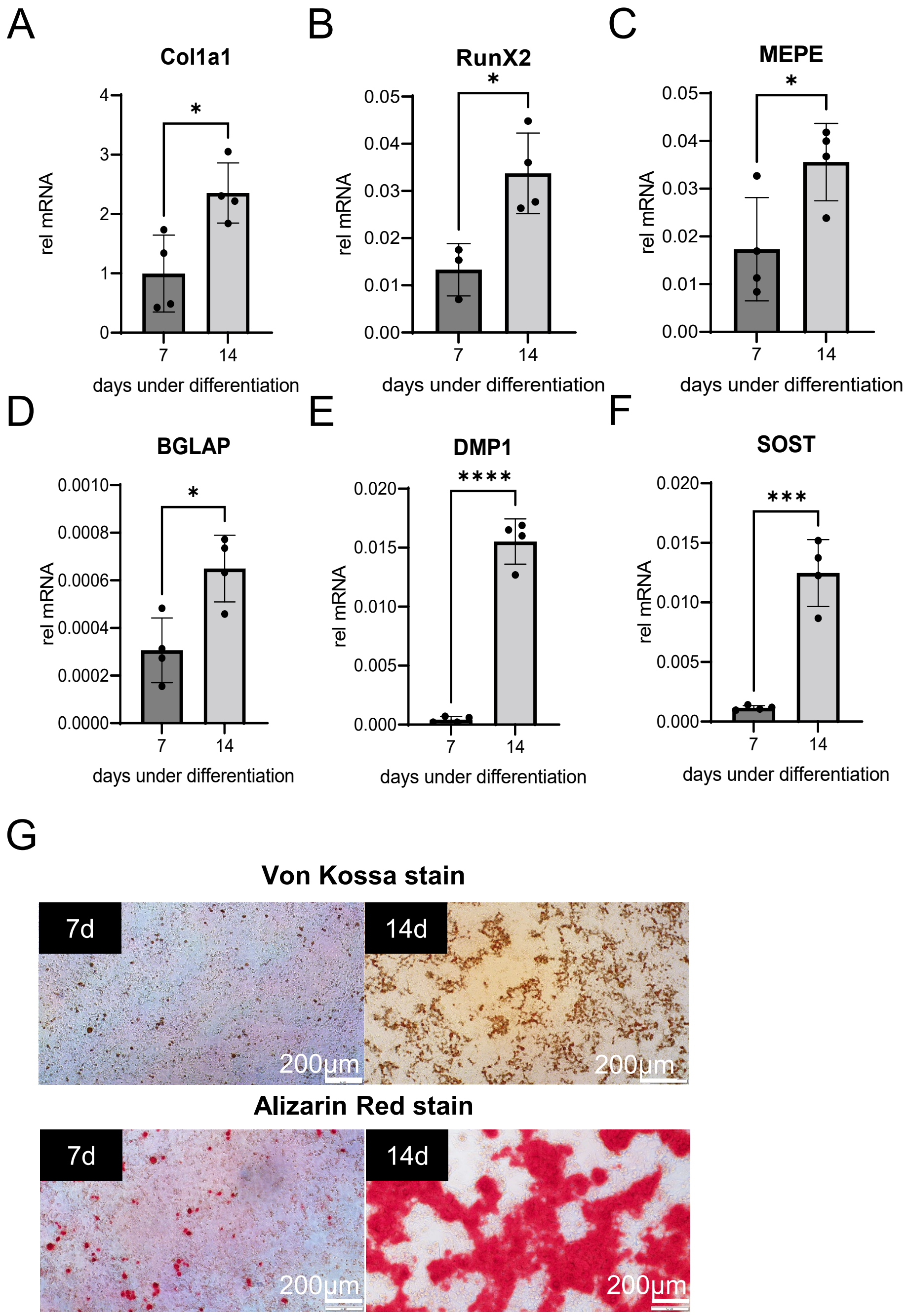


**Supplementary Figure 1:** Osteocytic differentiation of SaOS-2 cells. A-F) Osteogenic gene expression of osteocyte markers during differentiation relative to *18S* rRNA levels ($n$ = 4). *$p$ < 0.05, ***$p$ < 0.001 and ***$p$ < 0.0001. Gene expression was normalised to that of the housekeeping gene *18S* and calculated with the $2^{-\Delta Ct}$ method. G) Von Kossa stain for phosphate and Alizarin Red stain for calcium to observe mineralisation of osteocytes after 7 and 14 days of differentiation ($n$ = 4).
