## Supplementary material for "Antibiotics at Clinical Concentrations Show Limited Effectivity Against Acute and Chronic Intracellular *S. aureus* Infections in Osteocytes": Suppl Fig 2

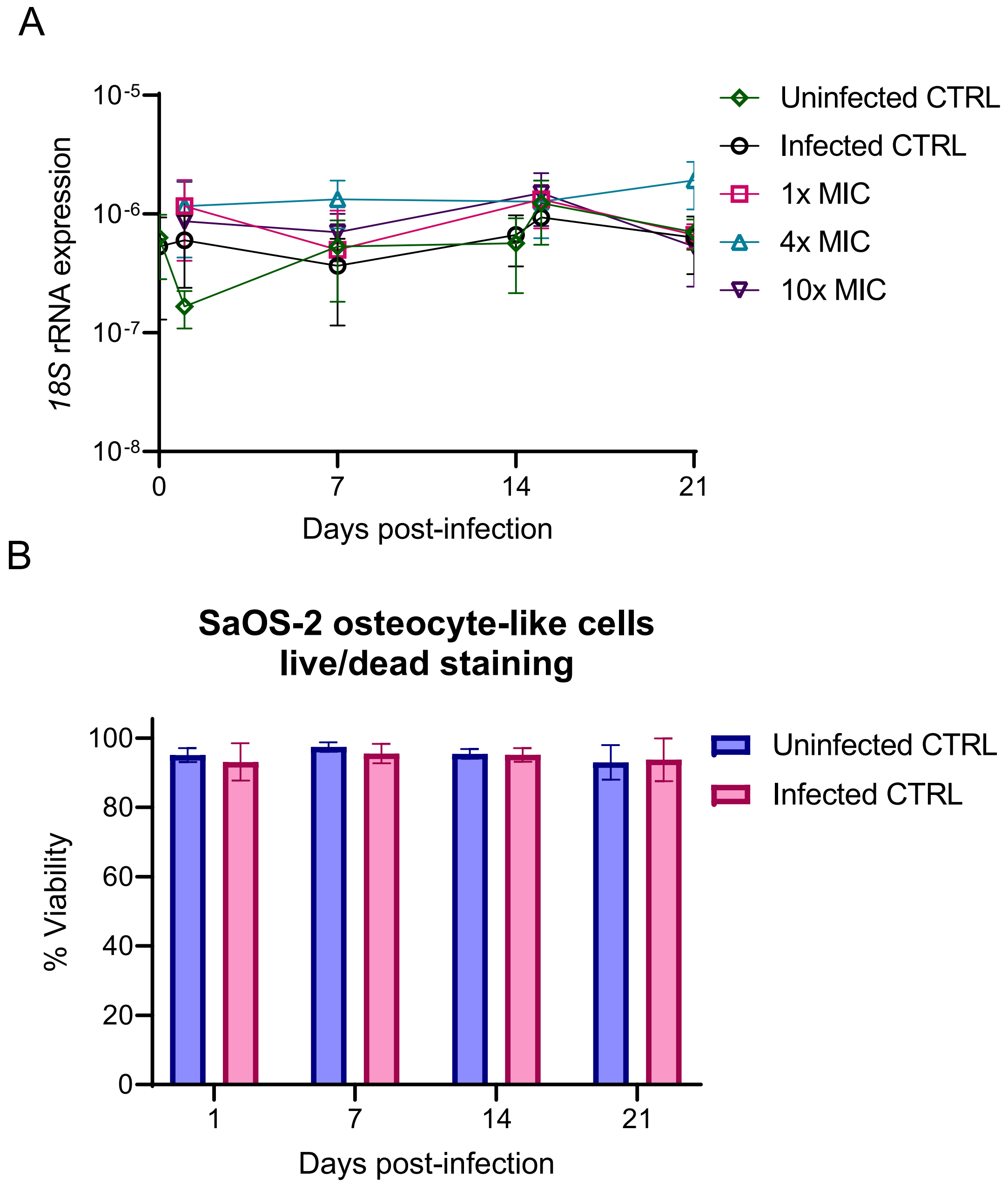


**Supplementary Figure 2:** Survival of host-cells during intracellular infection. A) Levels of *18S* rRNA over time in infected control, uninfected control or treatment with levofloxacin (as a representative example for all treatments; $n = 4$). B) Live/dead staining of infected and uninfected controls. Cells were cultured as described above, on Cell Imaging Plates (Eppendorf, Germany). Immediately before the infection or 1, 7, 15 and 21 days after the infection the media of infected and uninfected wells were incubated in the dark at 37°C/ 5% $CO_2$/ 1% $O_2$ with eBioscience TM Calcein Violet 450 AM Viability Dye (live, Invitrogen) and Ethidium Homodimer III (dead, Biotium) for 30 min. Confocal images were then taken with an Olympus FV3000 confocal microscope (Olympus, Tokyo, Japan) and processed with ImageJ software (Bethesda, USA) to obtain relative intensities of live/dead stained cells. Data shown as means ± standard error of the mean for 4 biological replicates, in at least 3 regions of interest per well.
